# Collaboration between two conserved sequence motifs drives ATPase stimulation of Hsp90 by Aha1

**DOI:** 10.1101/2025.06.10.658861

**Authors:** Desmond Prah Amoah, Rebecca Mercier, Gnin Alyousef, Jill L. Johnson, Paul LaPointe

## Abstract

Hsp90 is a dimeric molecular chaperone essential for the folding, stabilization, activation, and maturation of hundreds of client proteins, which are critical for cellular function. Co-chaperones, such as Aha1, play a key role in regulating the ATP-dependent Hsp90 client activation cycle by modulating Hsp90’s ATPase activity and controlling progression through the cycle. Two highly conserved motifs in Aha1—the NxNNWHW and RKxK motifs—are known to regulate specific aspects of the Hsp90 ATPase cycle. In this study, we demonstrate that the K60 residue within the RKxK motif facilitates the structural organization of the NxNNWHW motif prior to ATP hydrolysis. Mutation of the K60 residue partially impairs the *in vivo* functionality of yeast Aha1. Additionally, we reveal that each individual residue within the NxNNWHW motif modulates the ATPase rate and apparent affinity for ATP of Hsp90. These findings provide new insights into how conserved regions of Aha-type co-chaperones influence Hsp90 kinetics and its regulation of client protein folding.

## Introduction

The 90-kiloDalton heat shock protein 90 (Hsp90) is a dimeric molecular chaperone that facilitates the folding and maturation of a broad but specific group of substrates called client proteins (10; 48; 11; 51). These clients include membrane proteins, hormone receptors, transcription factors, and other key regulators of signal transduction, the cell cycle, and stress responses (54; 33; 7). The maturation of client proteins by the Hsp90 dimer takes place within an ATP-dependent functional cycle, during which Hsp90 undergoes extensive conformational changes involving both inter- and intra-protomer interactions (5; 8). The ATP-dependent functional cycle complex is regulated by an array of proteins called co-chaperones (6; 12; 18; 40; 47; 2; 21; 22). Co-chaperones influence ATP binding and hydrolysis, and modulate its interactions with client proteins (36; 46). Moreover, the Hsp90 functional cycle is controlled by post-translational modifications (PTMs), which can affect the Hsp90 conformational dynamics by altering client proteins and co-chaperone recruitment (30; 32; 57; 29; 38). Although the regulation of the Hsp90 functional cycle during client maturation is not well understood, it is evident that ATP hydrolysis is necessary for Hsp90 to efficiently mature clients (34; 35; 39).

Each Hsp90 monomer consists of three major domains: an N-terminal domain with an ATP-binding pocket, a middle domain (connected to the N domain by a long charged linker), and a C-terminal dimerization domain, which facilitates Hsp90 dimer formation (28; 41; 17). The interplay between these domains is essential for the catalytic activity of Hsp90 (8). The Hsp90 functional cycle begins with ATP binding to the N-terminal domain, which induces interactions of the γ phosphate of ATP with an Arg 380 (R380) residue in the middle domain (28; 9). This interaction leads to the docking of the N- and middle domains of Hsp90, initiating ATP hydrolysis (45). The ATP lid, a small region in the N-terminal domain, undergoes a conformational shift, trapping the ATP molecule and triggering the rotation of helices in the N-terminal domains of opposite monomers which exposes hydrophobic residues that facilitate N-terminal dimerization (8; 52). Although Hsp90 can adopt multiple structural states even in the absence of nucleotides (4), ATP binding is thought to lower the energy barrier between the apo (open) state and the closed, N-terminally dimerized state, enabling efficient client maturation (13).

The regulation of Hsp90 ATPase activity has become a focal point for understanding its role in client protein folding (36; 20). Aha1, Activator of Hsp90 ATPase 1, is the most robust known stimulator of the normally very low Hsp90 ATPase activity (25; 28). Aha1 is recruited to Hsp90 through various PTMs, such as phosphorylation and SUMOylation (30; 31; 29), and is crucial for the folding and activation of diverse proteins, including Glucocorticoid receptor, v-src and cystic fibrosis transmembrane conductance regulator (CFTR) (53; 14). Aha1 enhances the ATPase cycle of Hsp90, promoting its transition to a catalytically competent state, although the exact mechanism remains poorly understood (2; 22; 56; 26). Aha1 consists of two functional domains: an N-terminal domain and a C-terminal domain, connected by an unstructured linker (Figure 1a) (19; 44; 22; 56; 26). The N-terminal domain contains two highly conserved motifs—the NxNNWHW and RKxK motifs that are known to regulate Hsp90 ATPase activity (Figure 1a) (28; 15; 26; 16; 1). The NxNNWHW motif, located within the first 11 amino acids of yeast Aha1 and the first 27 amino acids of mammalian Aha1 (Ahsa1), is essential for maximal stimulation of the Hsp90 ATPase activity (26; 16; 1). Our previous work has shown that the NxNNWHW motif is required for the robust stimulation of the intrinsically low Hsp90 ATPase activity, and our recent findings indicate that this motif also enforces ordered ATP hydrolysis within the Hsp90 dimer (26; 16; 1). The *in vivo* action of Aha1 is also dependent on the presence of the NxNNWHW motif (26; 1). On the other hand, the role of the RKxK motif of Aha1 on the function of this co-chaperone, is not yet well understood.

**Figure 1.**
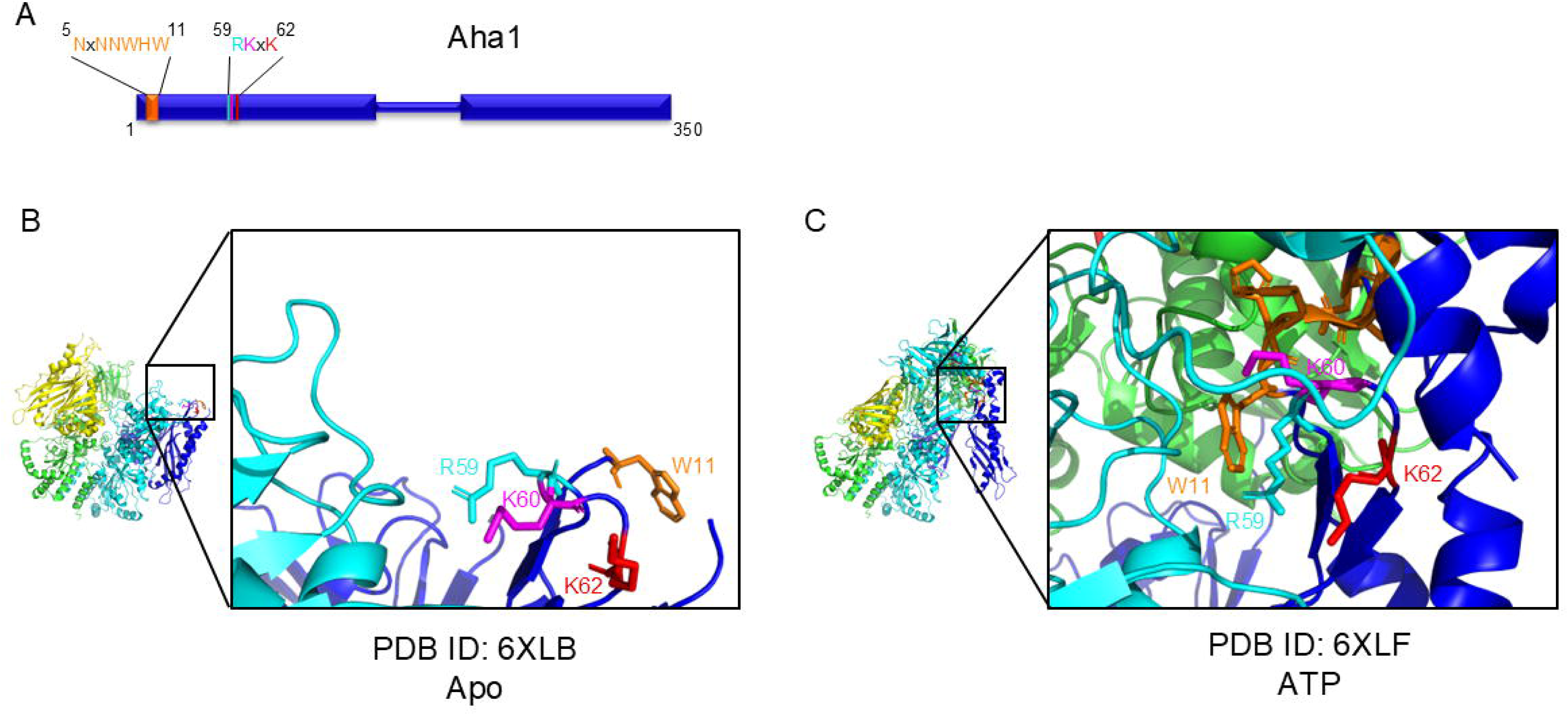
Interaction between the yeast Aha1 co-chaperone and Hsp90. (A) Aha1 is a co-chaperone of Hsp90 that consists of two distinct domains: an N domain and a C domain, connected by a flexible, unstructured linker. The N-terminal domain features two conserved motifs: the NxNNWHW motif and the RKxK motif. (B) The structure of the Aha1-Hsp90 complex in the nucleotide-free (Apo) state (PDB: 6XLB) reveals that both the N-terminal and C-terminal domains of Aha1 are bound to the middle domain of Hsp90. The RKxK motif is structured in this complex but the NxNNWHW motif (except for W11) is not. (C) In the nucleotide-bound state (PDB: 6XLF), the K60 residue within the RKxK motif of Aha1 (depicted in magenta) is repositioned towards the now-structured NxNNWHW motif (orange) in the N-terminal domain of Aha1. The R59 (cyan) and K62 (red) residues of the RKxK motif remain oriented towards the Hsp90 middle domain and away from the NxNNWHW motif (orange).

Recent cryo-EM structures have provided insights into how Aha1 interacts with Hsp90 to stimulate ATPase activity (24). In the absence of nucleotides, Aha1 binds to the middle domain of Hsp90 on opposing subunits, preventing the N-domains of Hsp90 from interacting with the middle domain (Figure 1b). Upon ATP binding, Aha1 undergoes a conformational change, allowing its N-terminal domain to interact with the N-domains of Hsp90. This leads to the structural ordering of the NxNNWHW motif, which then participates in interactions with various residues in both the N domains of Hsp90 and certain residues in the RKxK motif of the N domain of Aha1 (Figure 1c). Interestingly, the RKxK motif is situated beneath the NxNNWHW motif suggesting that they may act together to promote ATP hydrolysis.

To investigate the relationship between the conserved motifs in Aha1, we carried out site-directed mutagenesis of the NxNNWHW and RKxK motifs in yeast Aha1, substituting alanine for each residue. These alanine mutants were compared to wildtype Aha1 and Aha1^Δ11^ (Aha1 missing the first 11 amino acids containing the NxNNWHW motif) to assess their capacity to stimulate ATPase activity *in vitro*. We found that mutation of lysine 60 to alanine (K60A) in the RKxK motif significantly impaired ATPase stimulation by Aha1, mimicking the loss of the NxNNWHW motif. Furthermore, we showed that K60A mutation partially blocks the action of the NxNNWHW motif in yeast. Our results suggest that these conserved motifs act together to regulate the Hsp90 ATPase cycle.

## Online Methods

### Protein expression and Purification

*Saccharomyces cerevisiae* Hsc82, Hsc82^S25P^, Aha1, Aha1^Δ11^, Aha1^R59A^, Aha1^K60A^, Aha1^K62A^, Aha1^R59A/Δ11^, Aha1^K60A/Δ11^, Aha1^K62A/Δ11^, Aha1^N5A^, Aha1^N7A^, Aha1^N8A^, Aha1^N5/7/8A^, Aha1^W9A^, Aha1^H10A^, Aha1^W11A^, and Aha1^WHW/AAA^ were expressed in *Escherichia coli* strain BL21 (DE3) (New England Biolabs) from pET11d (Stratagene, La Jolla, CA, USA). Two versions of pET11d were used to express these proteins. Hsc82, and Hsc82^S25P^ were expressed with *N* terminal 6xHis tags and Aha1, Aha1^Δ11^, Aha1^R59A^, Aha1^K60A^, Aha1^K62A^, Aha1^R59A/Δ11^, Aha1^K60A/Δ11^, Aha1^K62A/Δ11^, Aha1^N5A^, Aha1^N7A^, Aha1^N8A^, Aha1^N5/7/8A^, Aha1^W9A^, Aha1^H10A^, Aha1^W11A^, and Aha1^WHW/AAA^ were expressed with C terminal 6xHis tags. Cells were grown at 37°C to an OD_600_ of 0.8-1.0 and induced with 1mM isopropyl-1-thio-D-galactopyranoside (IPTG). Cells expressing Hsc82, Hsc82^T22E^, Hsc82^V387E^, Hsc82^E33A^, Hsc82^D79N^, Hsc82^T22E/E33A^ and Hsc82^T22E/V387E^ were harvested after overnight growth at 16°C. Cells expressing Aha1 and Aha1^Δ11^ were harvested after 4hours at 37°C. Cells were harvested by centrifugation and stored at −80°C. Cells were resuspended in lysis buffer (50 mM KH_2_PO_4_, pH 8.0, 500 mM KCl, 10% Glycerol, 10 mM Imidazole, 5 mM β-mercaptoethanol) and lysed using Avestin Emulsiflex C3 (Avestin, Ottawa, Ontario, Canada).

Lysates were clarified by ultracentrifugation and His-tagged proteins were isolated on a HisTrap FF column using an AKTA Explorer FPLC (GE Healthcare). Isolated 6xHis-tagged proteins were then concentrated and subjected to Anion Exchange chromatography on a Hitrap Q FF column using an AKTA Explorer FPLC (GE Healthcare). The isolated proteins were further purified by size exclusion chromatography on a Hiload Superdex 200 pg column (GE Healthcare). Purity of each protein preparation was > 95% as verified by Coomassie-stained SDS-PAGE analysis.

### ATPase assays

ATPase assays were carried out using the enzyme coupled assay as previously described(2; 15; 56; 26; 55). All reactions were carried out in at least three independent experiments, with every condition in triplicate 100 µL (for 96-well plates) or 50 µL (for 384-well plates) reactions.

Absorbance at 340 nm was measured every 1 (for 96-well plates) or 2 (for 384-well plates) minutes for 90 min using a Bio Tek Synergy 4 and the path-length correction function. Average values of the experiments are shown with error expressed as standard error of the mean. The decrease in NADH absorbance at 340 nm was converted to micromoles of ATP using Beer’s Law and then expressed as a function of time^51^. The final conditions of all the reactions are 25 mM HEPES (pH 7.2), 12.5 or 16 mM NaCl (in titration and cycling experiments, respectively), 5 mM MgCl_2_, 1 mM DTT, 0.6 mM NADH, 2 mM ATP (co-chaperone titration and cycling experiments), 1 mM phosphoenol pyruvate (PEP), 2.5 µL of pyruvate kinase/lactate dehydrogenase (PK/LDH) (Sigma), and 5% DMSO. To correct for contaminating ATPase activity, identical reactions were quenched with 100 µM NVP-AUY922 and subtracted from unquenched reactions (DMSO control). In the titration experiments, 1 µM of Hsc82 or Hsc82^S25P^ was added to reactions containing either 1, 2, 4, 8 or 16 µM of Aha1 (or Aha1 mutants). The ATPase assay was started by the addition of the regenerating system consisting of MgCl_2_, DTT, NADH, ATP, PEP, PK/LDH. Fit lines were calculated according to the following equation (*Y* = ((*B*_max_\**X*)/(*K*_app_ + *X*)) + *X*_0_).

### Yeast growth assays

Yeast cells were transformed by lithium acetate methods and grown in either YPD or defined synthetic complete media supplemented with 2% dextrose. Growth was examined by spotting 10-fold serial dilutions of yeast cultures on appropriate media, followed by incubation for two days at the indicated temperature. Hsc82^S25P^ was expressed in strain JJ816 (*aha1hsc82hsp82*/Yep24-*HSP82*) after plasmid shuffling using 5-fluorootic acid (5-FOA) (Toronto Research Chemicals). Plasmids encoding wildtype and mutant Aha1 (p41KanTEF-Aha1-myc) have been described previously (26). G418 was obtained from Sigma. Quantification of yeast growth assays was done as previously described (37; 27).

## Results

### K60 is required for robust Hsp90 ATPase stimulation

The NxNNWHW and RKxK motifs in Aha1 are important for stimulating Hsp90 ATPase activity (28; 15; 26, 16, 1). However, while the NxNNWHW motif is necessary for robust ATPase stimulation by Aha1, the role of the RKxK motif is less well defined. To better understand the significance of the residues in the highly conserved RKxK motif, we introduced alanine residues at each position. We expressed and purified each of these constructs (harbouring a C-terminal 6xHis-tag) for testing in ATPase stimulation assays. The K60A mutation (Aha1^K60A^) caused a significant reduction in ATPase stimulation of Hsc82 (the constitutively-expressed yeast Hsp90) when compared to wildtype Aha1, Aha1^R59A^, or Aha1^K62A^ (Figure 2a/b/c). Interestingly, all three mutations reduced the apparent affinity of Aha1 for Hsc82 compared to wildtype with the greatest effect observed for Aha1^K60A^ (Figure 2d/e). This suggests that the conserved residues of RKxK motif play distinct roles in binding to Hsp90 and stimulation of ATPase activity.

**Figure 2.**
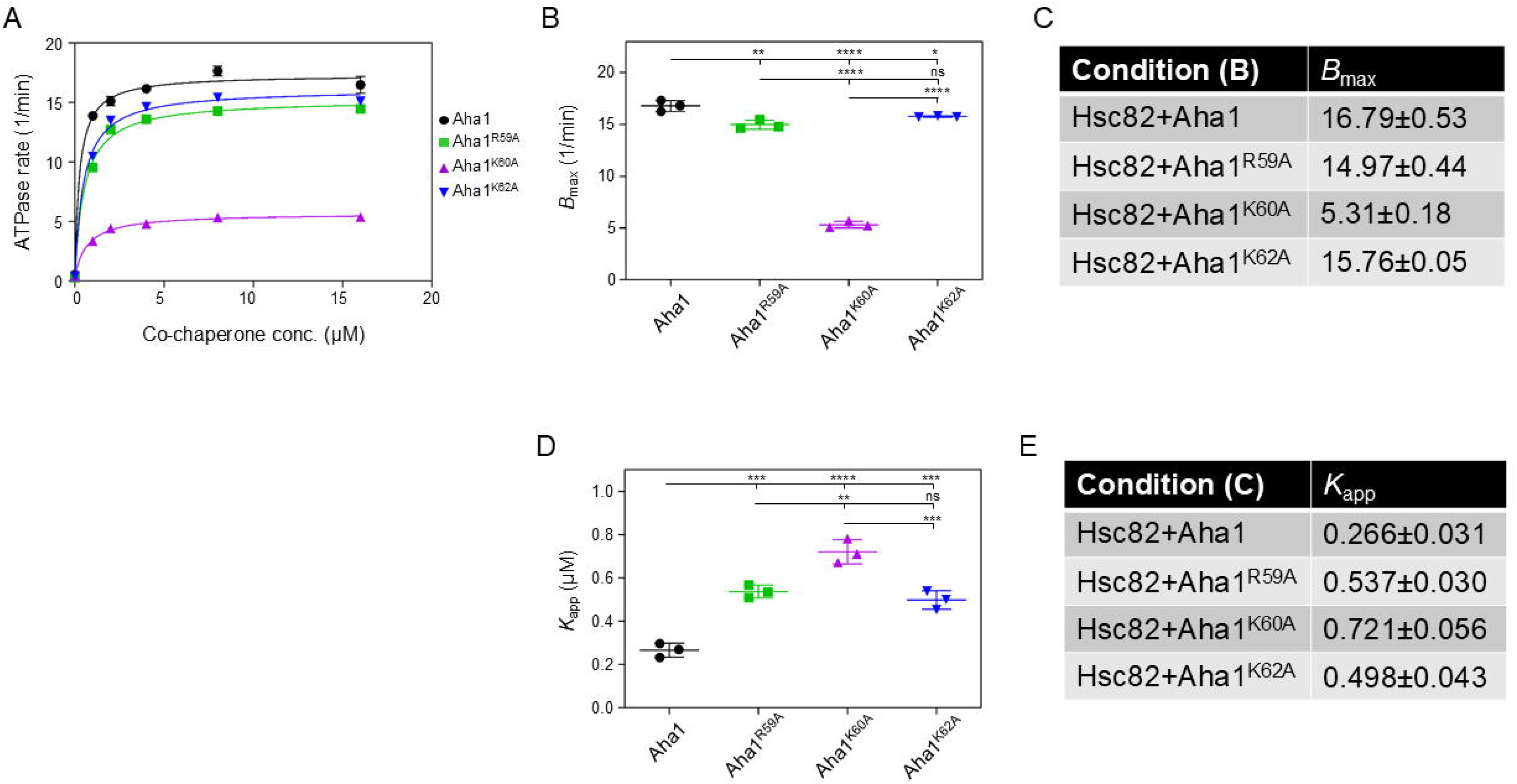
K60 is essential for strong ATPase stimulation by Aha1. (A) The stimulation of Hsc82 was measured using increasing concentrations of Aha1 (black circles), Aha1^R59A^ (green squares), Aha1^K60A^ (magenta triangles), and Aha1^K62A^ (inverted blue triangles). Each reaction contained 1 μM Hsc82 and varying concentrations of the co-chaperone (n = 3). Error bars represent the standard error of the mean. (B) The *B*_max_ values derived from the experiments in panel (A) are plotted. The *B*_max_ for Aha1^K60A^ was significantly lower than that for wildtype Aha1. In contrast, the *B*_max_ values for Aha1^R59A^ and Aha1^K62A^ were similar to that of wildtype Aha1. (C) Chart showing *B*^max^ values from (B). (D) *K*_app_ values derived from the experiments in panel (A) are plotted. The *K*_app_ for Aha1^K60A^ was significantly lower than that for wildtype Aha1. In contrast, the *K*_app_ values for Aha1^R59A^ and Aha1^K62A^ were similar to that of wildtype Aha1. (E) Chart showing *K*_app_ values from (D). Data Information: In A, data points are the mean of three independent triplicate experiments and error bars represent the standard error of the mean. Reactions contained 1μM Hsc82 and indicated concentration of co-chaperone (*N* = 3). Error bars in B and D represent the standard deviation. Statistical significance in B and D was determined using Tukey’s multiple comparisons test (\**p* < 0.05; \*\**p* < 0.01; \*\*\**p* < 0.001; \*\*\*\**p* < 0.0001).

### Mutation of K60 to an alanine mimics the loss of the NxNNWHW motif

As mentioned earlier, cryo-EM structures of the Hsp90-Aha1 complex show that the NxNNWHW motif is positioned near the RKxK residues upon ATP binding by Hsp90 (Figure 1b/c). Comparing the nucleotide-free and ATP-bound states reveals that K60, but not R59 and K62, reorients towards the NxNNWHW motif after ATP binding (Figure 1b/c). We hypothesized that the RKxK motif, and K60 in particular, enabled the participation of the NxNNWHW motif in ATPase stimulation. To test this, we measured the Aha1-mediated ATPase stimulation activity of single point mutations in the RKxK motif in combination with deletion of the NxNNWHW motif (Figure 3a-f). As expected, deletion of the NxNNWHW motif reduced ATPase stimulation by Aha1, Aha1^R59A^, and Aha1^K62A^ to the same degree. However, deletion of the NxNNWHW motif from Aha1^K60A^ did not result in a further decrease in ATPase stimulation (Figure 3c/d). In other words, the mutation of K60 to alanine had the same effect as deletion of the NxNNWHW motif and no significant additional reduction occurred when the two mutations are combined. That these two mutations mimic one another suggested to us that K60 is required for the action of the NxNNWHW motif during ATPase stimulation.

**Figure 3.**
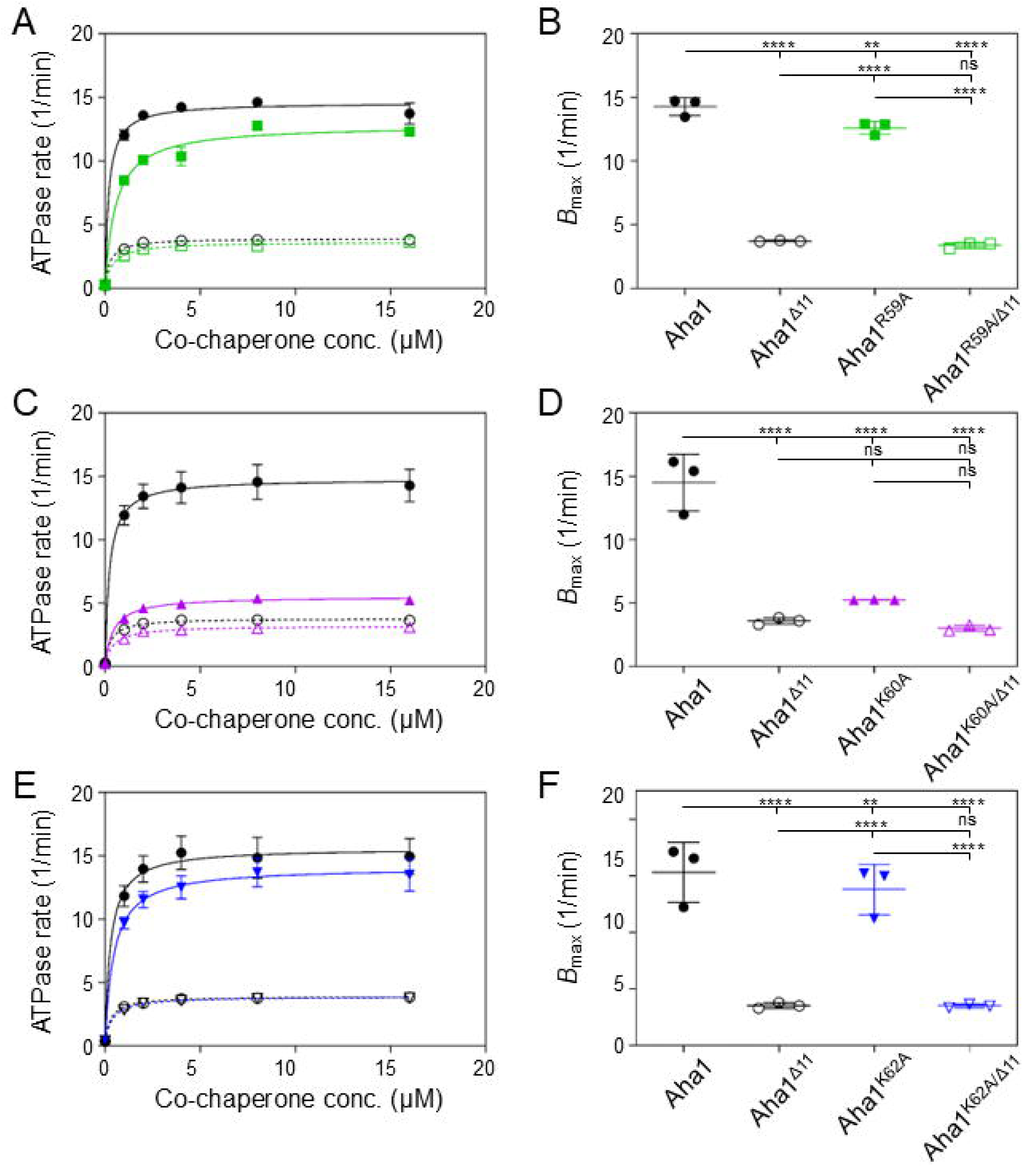
The K60A mutation mimics the loss of the NxNNWHW motif. (A) Stimulation of Hsc82 by increasing concentrations of Aha1 (black circles), Aha1^Δ11^ (open black circles), Aha1^R59A^ (green circles) or Aha1^R59A/Δ11^ (open green circles). (B) The *B*_max_ values derived from the experiments in panel (A) are plotted. (C) Stimulation of Hsc82 by increasing concentrations of Aha1 (black circles), Aha1^Δ11^ (open black circles), Aha1^K60A^ (purple triangles) or Aha1^K60A/Δ11^ (open purple triangles). (D) The *B*_max_ values derived from the experiments in panel (C) are plotted. (E) Stimulation of Hsc82 by increasing concentrations of Aha1 (black circles), Aha1^Δ11^ (open black circles), Aha1^K62A^ (blue inverted triangles) or Aha1^K62A/Δ11^ (open blue inverted triangles). (F) The *B*_max_ values derived from the experiments in panel (E) are plotted. Data Information: In A, C, and E, data points are the mean of three independent triplicate experiments and error bars represent the standard error of the mean. Reactions contained 1μM Hsc82 and indicated concentration of co-chaperone (*N* = 3). Error bars in B, D, and F represent the standard deviation. Statistical significance in B, D, and F was determined using Tukey’s multiple comparisons test (\*\**p* < 0.01; \*\*\*\**p* < 0.0001).

### Mutation of K60 to an alanine has the same effects on the apparent affinity for ATP as deletion of the NxNNWHW motif

In previous studies, we demonstrated that the loss of the NxNNWHW in yeast Aha1 affects the apparent affinity of Hsp90 for ATP (26). Since the K60A mutation blocks the action of the NxNNWHW motif, we hypothesized that K60A would increase the apparent affinity of Hsp90 for ATP similar to the effects observed for the deletion of the NxNNWHW motif. Consistent with our hypothesis, the apparent affinity of Hsc82 for ATP in the presence of Aha1^K60A^ was increased to the same degree as for Aha1^Δ11^ (Figure 4a/b). Importantly, there were no significant difference in apparent affinity of ATP for Hsc82 in the presence of wildtype Aha1, Aha1^R59A^, or Aha1^K62A^ (Figure 4a/b). This further supported the idea that K60 is required for the action of the NxNNWHW motif.

**Figure 4.**
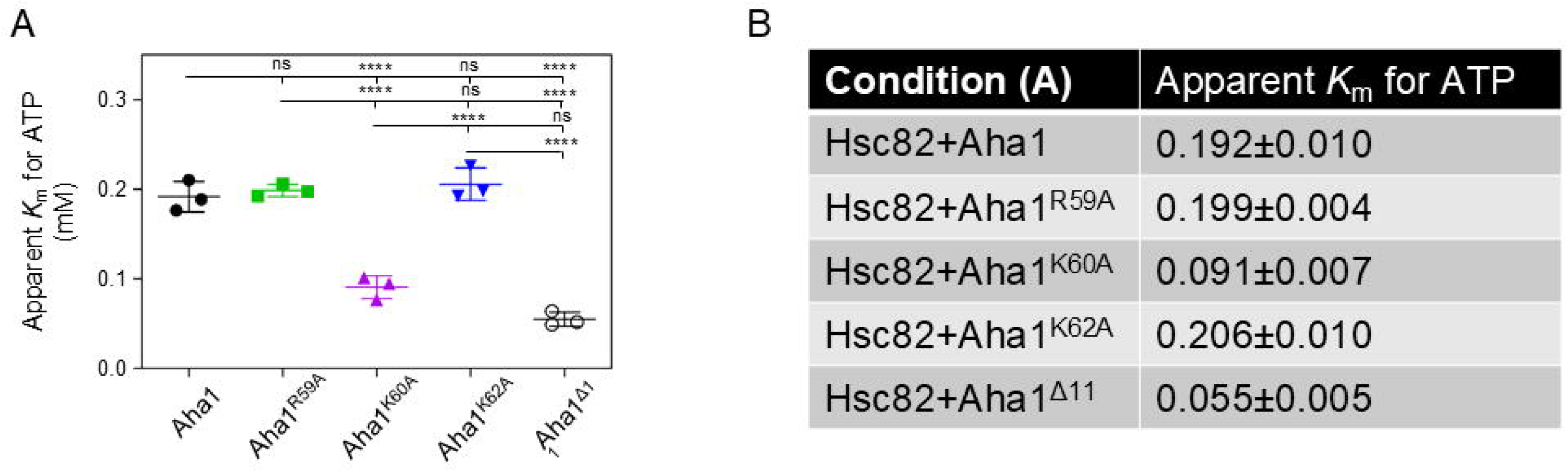
The K60A mutation has the same effect on the apparent affinity for ATP as deletion of the NxNNWHW motif. (A) The apparent Km for ATP of Hsc82 was measured in the presence of Aha1 (black circles), Aha1^R59A^ (green squares), Aha1^K60A^ (purple triangles) Aha1^K62A^ (blue inverted triangles), and Aha1^Δ11^ (open black circles). ATPase activity was measured using increasing ATP concentrations (12.5, 25, 50, 100, 200, 400, 800, 1600 μM), and the resulting ATPase rates were analyzed using the Michaelis-Menten non-linear regression function in GraphPad Prism. The curve fits all had *R*^2^ values >0.9. The apparent *K*_m_ values for three independent experiments are plotted and the error bars represent the standard error (n=3). Statistical significance was calculated using a Tukey’s multiple comparisons test (\*\*\*\**p* < 0.0001). (B) Table showing *K*_m_ values plotted in (A).

### K60A mutation blocks the action of the NxNNWHW of Aha1 in vivo

We previously showed that the NxNNWHW motif is required for the *in vivo* action of Aha1 in yeast (26). Specifically, overexpression of Aha1, but not Aha1^Δ11^, rescued the temperature-sensitive growth phenotype exhibited by yeast expressing Hsc82^S25P^. Since K60A mutation mimics deletion of the NxNNWHW motif, we hypothesized that the K60A mutation would prevent the rescue of temperature sensitive growth of yeast expressing Hsc82^S25P^. To test this, we expressed our different Aha1 constructs in a yeast strain expressing Hsc82^S25P^ in which the endogenous *AHA1* gene had been deleted. This strain grew poorly at elevated temperatures but was restored with the overexpression of Aha1, Aha1^R59A^, and Aha1^K62A^, but not Aha1^K60A^ (Figure 5a). Importantly, all of our Aha1 constructs were expressed to a comparable degree suggesting that Aha1^K60A^ is functionally impaired in a manner that Aha1^R59A^ and Aha1^K62A^ are not (Figure 5b).

**Figure 5.**
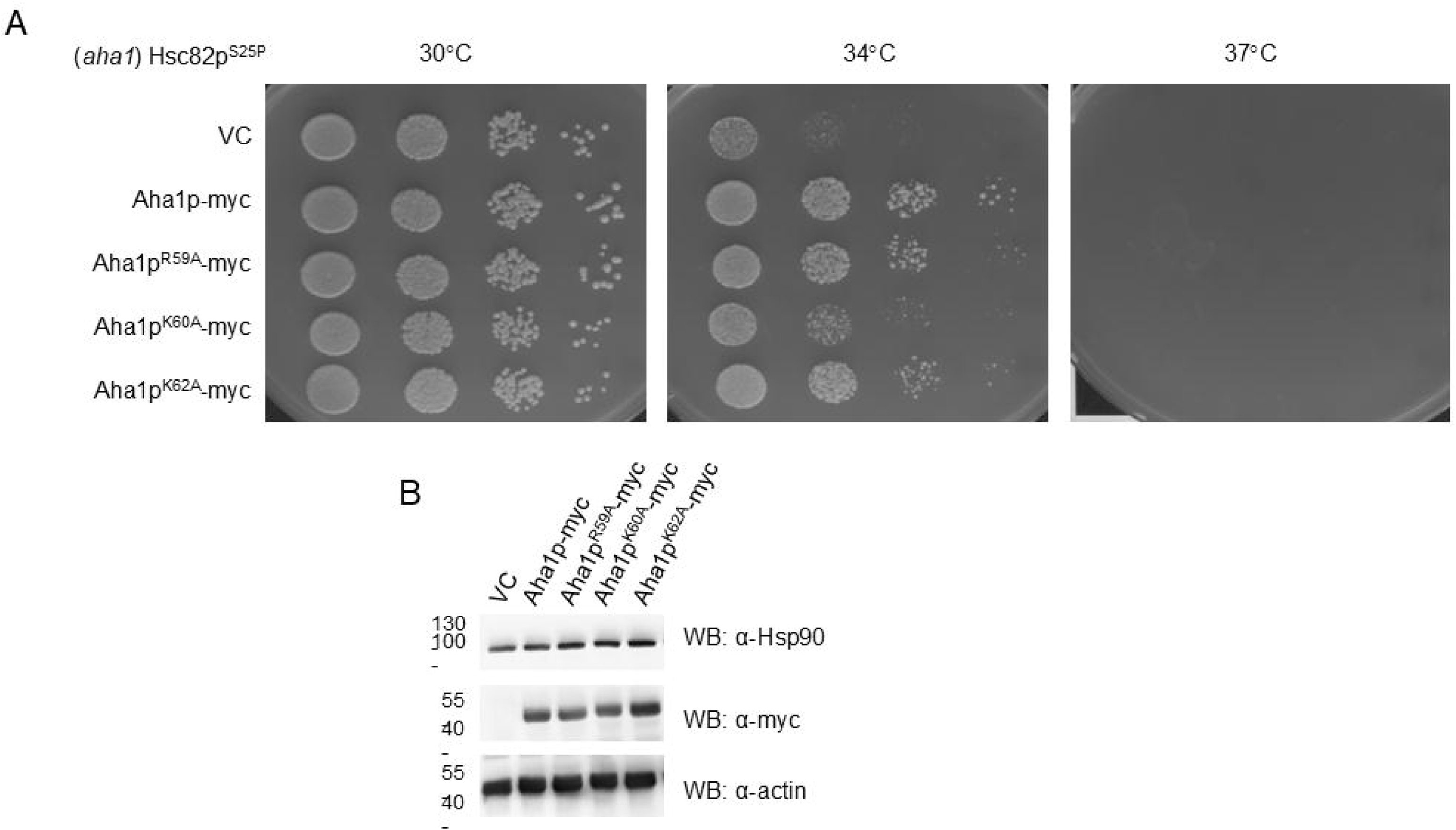
K60A mutation impairs the action of the NxNNWHW of Aha1 in yeast. (A) Yeast strains expressing Hsc82^S25P^ as the sole source of Hsp90 exhibit temperatures sensitive growth. Yeast cells containing plasmids encoding the specified C-terminally myc-tagged Aha1 co-chaperones were cultured overnight at 30°C in YPD medium supplemented with 200 mg/L Geneticin (for Aha1 plasmid selection). The cultures were then diluted to a concentration of 1 × 10^8 cells per milliliter. Ten-fold serial dilutions were prepared, and 10 μL aliquots were spotted onto YPD agar plates containing 200 mg/L Geneticin. The plates were incubated for 2 days at temperatures of 30°C, 34°C, or 37°C. Rescue of growth of Hsc82^S25P^ yeast by Aha1^K60A^ was impaired compared to wildtype Aha1, Aha1^R59A^, and Aha1^K62A^. (B) Western blot of total lysates extracted from the yeast strains shown in (A) were probed with anti-Hsp90, anti-actin, and anti-myc (for Aha1) antibodies.

### ATPase stimulation of Hsc82^S25P^ by RKxK mutants

We previously showed that ATPase stimulation of Hsc82^S25P^ by Aha1 requires the NxNNWHW motif (26). Importantly, deletion of the NxNNWHW motif and mutation of K60 (but not of R59 and K62) both resulted in a near complete loss of ATPase stimulation of Hsc82^S25P^ (Figure 6a/b) suggesting that these two alterations result in the same impairment.

**Figure 6.**
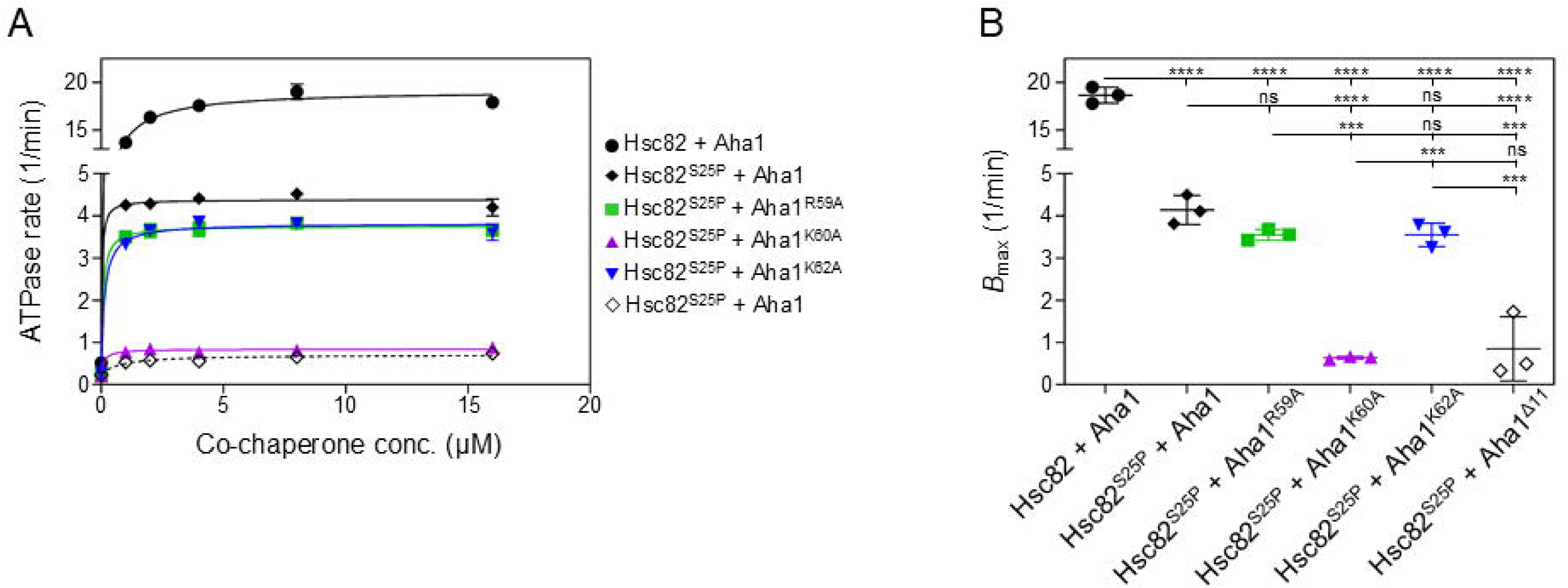
The stimulation of Hsc82^S25P^ ATPase activity by Aha1 depends on the K60 residue, which supports the function of the NxNNWHW motif. (A) The ATPase activity of both wildtype Hsc82 (black circles) and Hsc82^S25P^ (black diamonds) was enhanced by increasing concentrations of Aha1. Hsc82^S25P^ ATPase activity was stimulated by Aha1^R59A^ (green squares) and Aha1^K62A^ (blue inverted triangles), but not by Aha1^K60A^ (purple triangles) or Aha1^Δ11^ (open black diamonds). (B) The *B*_max_ values derived from the experiments in panel (A) are plotted. Data Information: In A, data points are the mean of three independent triplicate experiments and error bars represent the standard error of the mean. Reactions contained 1μM Hsc82 and indicated concentration of co-chaperone (*N* = 3). Error bars in B represent the standard deviation. Statistical significance in B was determined using Tukey’s multiple comparisons test (\*\*\**p* < 0.001; \*\*\*\**p* < 0.0001).

### All residues in the NxNNWHW motif contribute to its function

The cryo-EM structure of the Hsp90-Aha1 complex in the ATP-bound state shows that the NxNN and the WHW portion of the motif are positioned on opposite sides of a turn in the polypeptide backbone (Figure 7a). More specifically, the WHW portion of the motif appear to make contact with the RKxK motif and parts of Hsp90 while most of the NxNN portion makes contact with other parts of Aha1. To determine the contribution of each residue to ATPase stimulation we constructed Aha1 point mutants at each position as well as two different combination mutants where all asparagine residues of the NxNN were mutated or all residues of the WHW were mutated. Surprisingly, all mutants of Aha1 were profoundly compromised in their ability to stimulate the ATPase activity of Hsc82 (Figure 7b/c). Only Aha1^N7A^ could stimulate the ATPase activity of Hsc82 to a degree greater than Aha1 lacking the entire NxNNWHW motif (Aha1^Δ11^). This suggests that each residue in the NxNNWHW motif is required for the robust Aha1-mediated ATPase stimulation of Hsp90.

**Figure 7.**
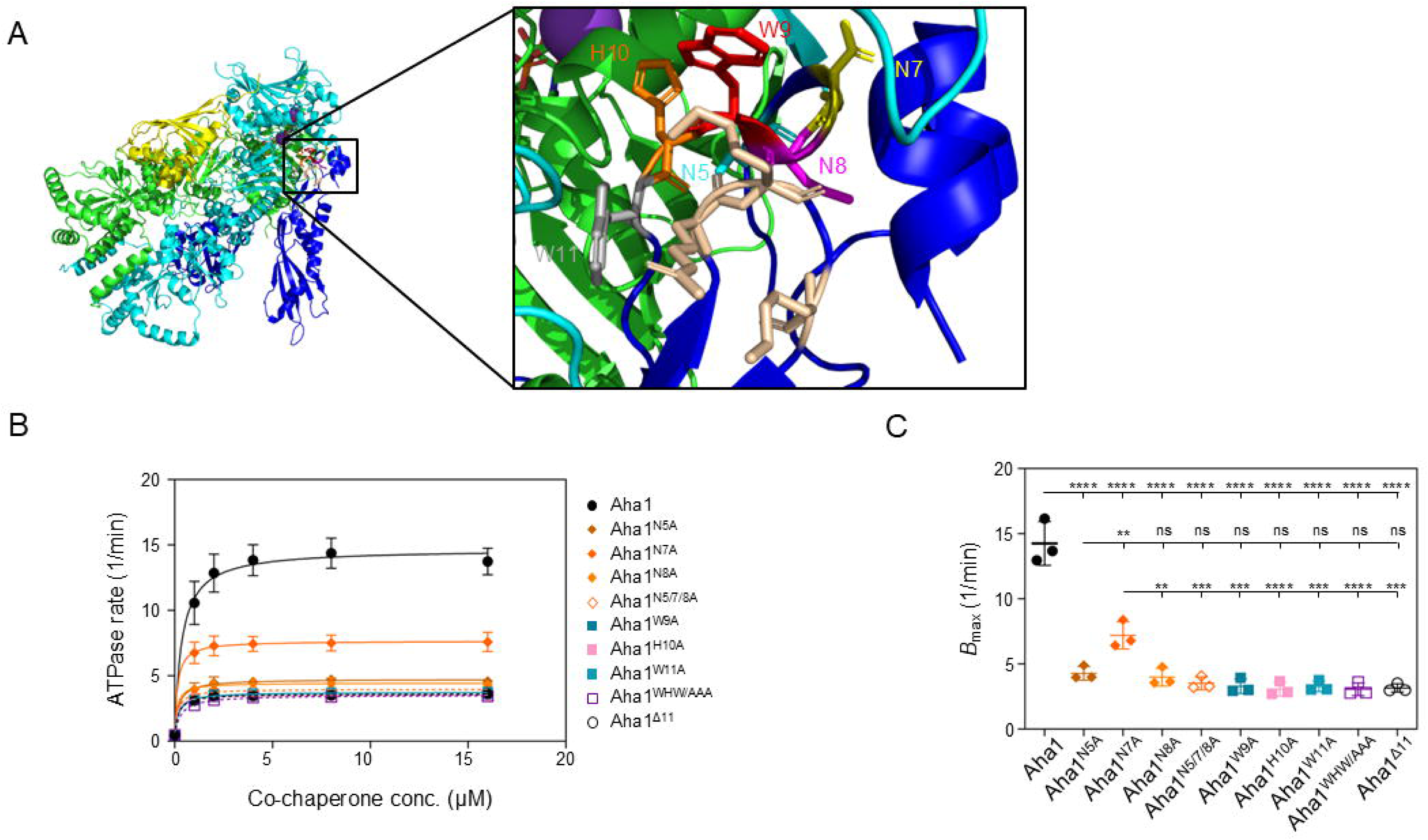
Mutation of each residue of the NxNNWHW motif to alanine mimics deletion of NxNNWHW motif. (A) Schematic of Hsp90-Aha1 complex in the nucleotide-bound state (PDBID: 6XLF) depicting the N5 (cyan), N7 (yellow), N8 (magenta), W9 (red), H10 (orange) and W11(grey) residues of the NxNNWHW motif located in the N-domain of Aha1. RKxK motif is shown in tan. (B) Stimulation of Hsc82 ATPase activity by increasing concentrations of Aha1 (black circles), N5A, N7A, N8A (each in a different shaded of orange diamond), N5A/N7A/N8A (open orange diamonds), W9A (dark turquoise squares), H10A (pink squares), W11A (light turquoise squares), W9A/H10A/W11A (open purple squares) and Aha1^Δ11^ (open black circles). (C) The *B max* values derived from the experiments in panel (B) are plotted. Data Information: In B, data points are the mean of three independent triplicate experiments and error bars represent the standard error of the mean. Reactions contained 1μM Hsc82 and indicated concentration of co-chaperone (*N* = 3). Error bars in C represent the standard deviation. Statistical significance in C was determined using Tukey’s multiple comparisons test (\*\**p* < 0.001; \*\*\**p* < 0.001; \*\*\*\**p* < 0.0001).

### Mutation of the NxNNWHW motif to alanine increases the apparent for ATP

We previously demonstrated that deletion of the NxNNWHW motif of Aha1 increases the apparent affinity of Hsp90 for ATP (26). To determine if the point mutations we introduced into the NxNNWHW motif also corresponded to a decrease in the apparent *K*_m_ for ATP, we tested our constructs in ATPase stimulation assays with increasing concentrations of ATP. Every mutation in the NxNNWHW (except N7) increased the apparent affinity of Hsc82 for ATP compared to wildtype Aha1 and to a similar extent as Aha1^Δ11^ (Figure 8a-d). That the N7A mutation resulted in the smallest change in maximal stimulation rate and apparent affinity for ATP suggests that this residue, despite being strongly conserved, is the least important for the function of the NxNNWHW motif.

**Figure 8.**
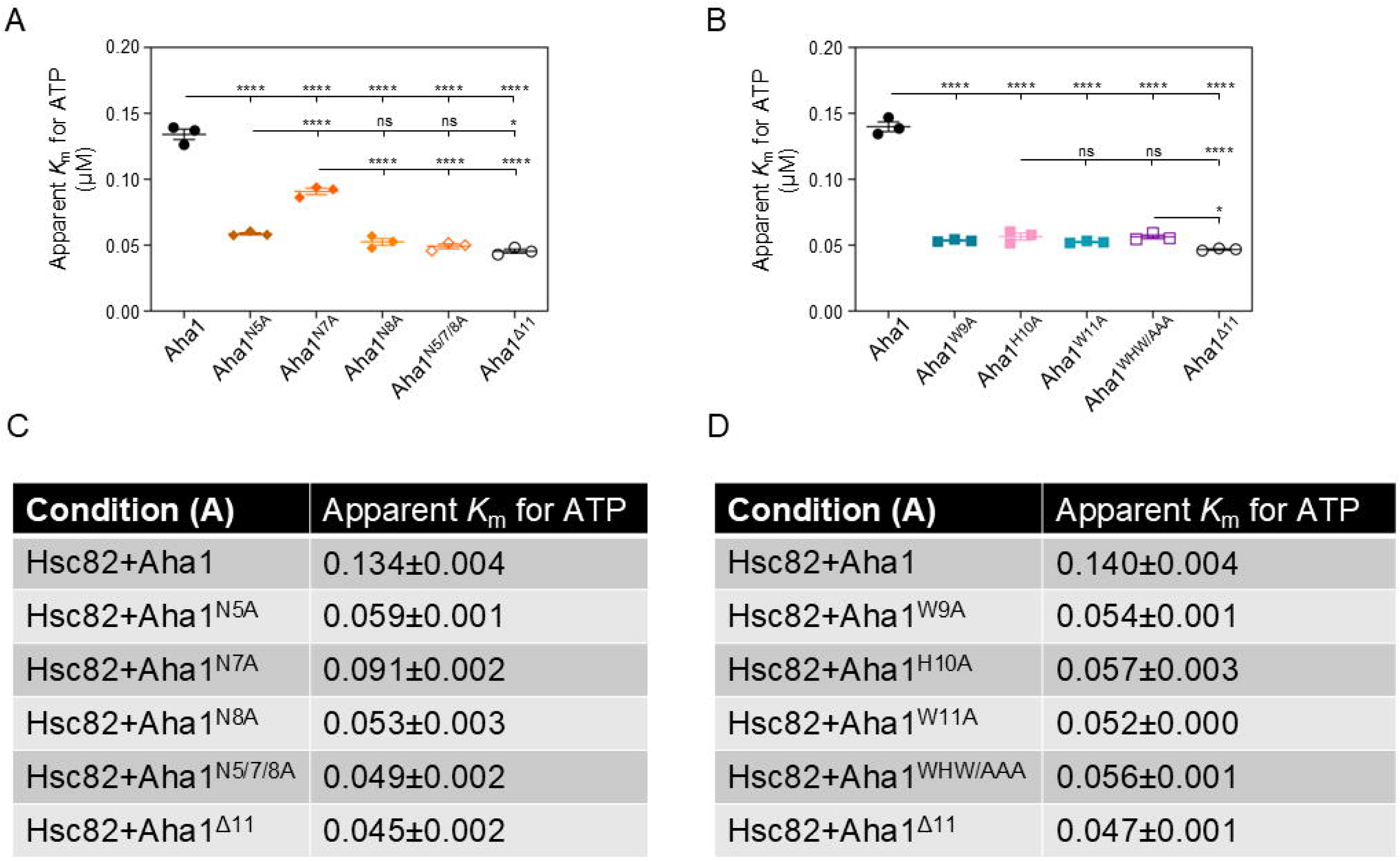
Mutations in the NxNNWHW motif increases the apparent for ATP of Hsc82. (A) The apparent *K*_m_ for ATP of Hsc82 was measured in the presence of Aha1 (black circles), N5A, N7A, N8A (each in a different shaded of orange diamond), N5A/N7A/N8A (open orange diamonds), and Aha1^Δ11^ (open black circles). (B) The apparent *K*_m_ for ATP of Hsc82 was measured in the presence of Aha1 (black circles), W9A (dark turquoise squares), H10A (pink squares), W11A (light turquoise squares), W9A/H10A/W11A (open purple squares), and Aha1^Δ11^ (open black circles). ATPase activity was measured using increasing ATP concentrations (12.5, 25, 50, 100, 200, 400, 800, 1600 μM), and the resulting ATPase rates were analyzed using the Michaelis-Menten non-linear regression function in GraphPad Prism. The curve fits all had *R*^2^ values >0.9. The apparent *K*_m_ values for three independent experiments are plotted and the error bars represent the standard error (n=3). Statistical significance was calculated using a Tukey’s multiple comparisons test (\**p* < 0.05; \*\*\*\**p* < 0.0001). (C) Table showing *K*_m_ values plotted in (A). (D) Table showing *K*_m_ values plotted in (B).

## Discussion

Cryo-EM structures of Aha1 in complex with Hsp90 have provided insight into the basis for ATPase regulation of Hsp90 (24). In the nucleotide-free state, the N and C-terminal domains of Aha1 are each bound to the middle domain of a different subunit of the Hsp90 dimer. The R59, K60 and K62 residues of the RKxK motif are positioned in the vicinity of the middle domain residues (310-390) of Hsp90 (Figure 1b). Upon ATP binding by Hsp90, the Aha1 N domain tilts towards the now-resolved N domains of Hsp90 and the NxNNWHW motif acquires structure near residues in Hsp90 that are important for ATP hydrolysis. Interestingly, K60 reorients towards the now-structured NxNNWHW motif within 4 angstroms of W9 and W11 (Figure 1c). That the K60A mutation mimics the deletion of the NxNNWHW in all the assays we tested suggests that this residue is required for the action of the NxNNWHW motif. Moreover, the function of the NxNNWHW motif is sensitive to mutations at any amino acid position suggesting an intricate network of interactions that lead to the hydrolysis of ATP as well as the reverse process where ADP is released (26).

Interestingly, despite K60 being the only residue that impaired the function of the NxNNWHW motif (Figure 2a-c), all three residues of RKxK are important for binding to Hsp90 (Figure 9). Mutations of any of the three conserved residues resulted in a significant reduction in apparent binding affinity (Figure 2d, e). Furthermore, the mutation of K60, but not R59 or K62, impacted the affinity of Hsp90 for ATP, consistent with the effects observed from the deletion of the NxNNWHW motif (Figure 4a, b). This suggests that the K60 residue of the RKxK motif of Aha1 plays a crucial role in controlling whether Hsp90 binds ATP tightly or loosely, as well as in how ATP is retained in the nucleotide-binding pocket through regulation of lid dynamics which might be through supporting the NxNNWHW motif. This would be entirely consistent with work suggesting that nucleotide exchange (and not hydrolysis) is the only absolute requirement for Hsp90 function in cells (58; 43; 42). We tested the biological activity of Aha1 in yeast expressing Hsc82^S25P^. Serine 25 has previously been shown to be a key phosphorylation site required for the induction of autophagy (3). That Aha1 overexpression rescues the growth of yeast expressing Hsc82^S25P^ likely suggests that larger structural perturbations caused by the introduction of a proline at this site can be corrected by proper assemble of the NxNNWHW and RKxK motifs upon Aha1 binding.

**Figure 9.**
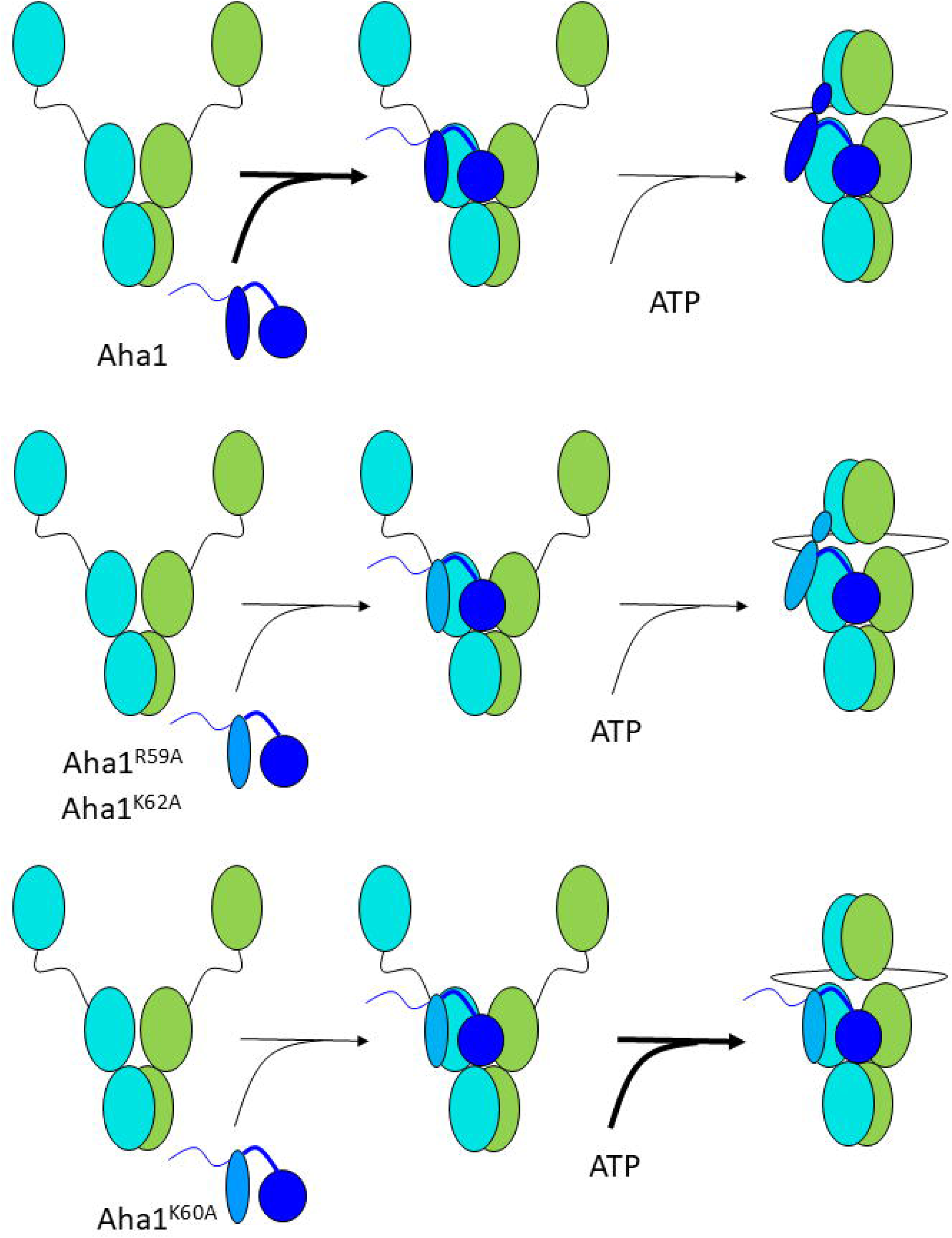
Schematic showing a model for the role of the RKxK motif in Hsp90 ATPase stimulation. Top panel; Aha1 binds to Hsp90 and stimulates ATPase activity. Middle panel; Mutation of either R59 or K62 impair binding affinity to Hsp90 but not ATPase stimulation. Bottom panel; Mutation of K60 impairs binding affinity to Hsp90 and ATPase stimulation by preventing the action of the NxNNWHW motif.

Our data indicate that each residue in the NxNNWHW motif in yeast Aha1 (except N7) plays a similar role in regulating ATPase activity and the apparent affinity of Hsp90 for ATP (Figure 7/8). We previously showed that ATPase stimulation by mammalian Aha1 (Ahsa1) was similarly impacted by mutations in the NxNN and WHW sections of the motif (16). In contrast, only mutations in the WHW portion of the NxNNWHW motif of mammalian Ahsa1 affected the apparent affinity for ATP. This suggests that the yeast and mammalian Aha1 proteins regulate Hsp90 differently despite the complete conservation of this sequence motif. A possible explanation for this discrepancy is that yeast Aha1 lacks the first 20 amino acids found in other Aha1 from other non-yeast organisms (16). This 20 amino acid N-terminal extension may interact with clients directly (50; 23; 49), with Hsp90 directly, or with other parts of Ahsa1. Without structural information regarding the human Ahsa1-Hsp90 complex, it is difficult to determine how subtle nucleotide-dependent conformational changes occur.

The data presented here suggest that the complete RKxK motif of Aha1 plays a critical role in binding Aha1 to Hsp90, perhaps following recruitment through modifications on specific Hsp90 residues (31; 32, 57, 29). Specifically, the K60 residue is likely essential for the structural alignment of the NxNNWHW motif of Aha1, positioning it near key catalytic residues that are necessary for ATP hydrolysis. Additionally, K60 may be crucial in regulating ATP binding within the ATP-binding pocket by influencing lid dynamics, which in turn impacts ATP-dependent client processing. While our findings provide valuable insights into how the different components of Aha1’s conserved motifs influence the Hsp90 chaperone cycle, further research is needed to fully understand how the RKxK motif and NxNNWHW specifically contributes to the regulation of client protein folding.

## Acknowledgments

This work was supported with funding from the Canadian Institutes of Health Research (178282) and the Natural Sciences and Engineering Research Council (RGPIN 2019 04967) (P.L.), as well as the National Institutes of Health (NIH R01 GM127675) (J.L.J.).

## Conflict of Interest Statement

The authors declare that they have no conflict of interest.

## Notes

### Competing Interest Statement

The authors have declared no competing interest.

### Summary of Updates

The methods section was mistakenly omitted from the previous version. This revision now includes the methods section.

## References

1. Amoah DP, Hussein SK, Johnson JL, LaPointe P (2025) Ordered ATP hydrolysis in the Hsp90 chaperone is regulated by Aha1 and a conserved post-translational modification. Protein Sci 34:e5255. PMID: 39665290 {Medline}

2. Armstrong H, Wolmarans A, Mercier R, Mai B, LaPointe P (2012) The co-chaperone Hch1 regulates Hsp90 function differently than its homologue Aha1 and confers sensitivity to yeast to the Hsp90 inhibitor NVP-AUY922. PLoS One 7:e49322. PMID: 23166640 {Medline}

3. Backe SJ, Sager RA, Heritz JA, Wengert LA, Meluni KA, Aran-Guiu X, Panaretou B, Woodford MR, Prodromou C, Bourboulia D, Mollapour M (2023) Activation of autophagy depends on Atg1/Ulk1-mediated phosphorylation and inhibition of the Hsp90 chaperone machinery. Cell Rep 42:112807. PMID: 37453059 {Medline}

4. Bron P, Giudice E, Rolland JP, Buey RM, Barbier P, Diaz JF, Peyrot V, Thomas D, Garnier C (2008) Apo-Hsp90 coexists in two open conformational states in solution. Biol Cell 100:413–425. PMID: 18215117 {Medline}

5. Chadli A, Bouhouche I, Sullivan W, Stensgard B, McMahon N, Catelli MG, Toft DO (2000) Dimerization and N-terminal domain proximity underlie the function of the molecular chaperone heat shock protein 90. Proc Natl Acad Sci U S A 97:12524–12529. PMID: 11050175 {Medline}

6. Chang HC, Nathan DF, Lindquist S (1997) In vivo analysis of the Hsp90 cochaperone Sti1 (p60). Mol Cell Biol 17:318–325. PMID: 8972212 {Medline}

7. Citri A, Gan J, Mosesson Y, Vereb G, Szollosi J, Yarden Y (2004) Hsp90 restrains ErbB-2/HER2 signalling by limiting heterodimer formation. EMBO Rep 5:1165–1170. PMID: 15568014 {Medline}

8. Cunningham CN, Krukenberg KA, Agard DA (2008) Intra- and intermonomer interactions are required to synergistically facilitate ATP hydrolysis in Hsp90. J Biol Chem 283:21170–21178. PMID: 18492664 {Medline}

9. Cunningham CN, Southworth DR, Krukenberg KA, Agard DA (2012) The conserved arginine 380 of Hsp90 is not a catalytic residue, but stabilizes the closed conformation required for ATP hydrolysis. Protein Sci 21:1162–1171. PMID: 22653663 {Medline}

10. Echeverria PC, Bernthaler A, Dupuis P, Mayer B, Picard D (2011) An interaction network predicted from public data as a discovery tool: application to the Hsp90 molecular chaperone machine. PLoS One 6:e26044. PMID: 22022502 {Medline}

11. Eckl JM, Richter K (2013) Functions of the Hsp90 chaperone system: lifting client proteins to new heights. Int J Biochem Mol Biol 4:157–165. PMID: 24380020 {Medline}

12. Fang Y, Fliss AE, Rao J, Caplan AJ (1998) SBA1 encodes a yeast hsp90 cochaperone that is homologous to vertebrate p23 proteins. Mol Cell Biol 18:3727–3734. PMID: 9632755 {Medline}

13. Hessling M, Richter K, Buchner J (2009) Dissection of the ATP-induced conformational cycle of the molecular chaperone Hsp90. Nat Struct Mol Biol 16:287–293. PMID: 19234467 {Medline}

14. Holmes JL, Sharp SY, Hobbs S, Workman P (2008) Silencing of HSP90 cochaperone AHA1 expression decreases client protein activation and increases cellular sensitivity to the HSP90 inhibitor 17-allylamino-17-demethoxygeldanamycin. Cancer Res 68:1188–1197. PMID: 18281495 {Medline}

15. Horvat NK, Armstrong H, Lee BL, Mercier R, Wolmarans A, Knowles J, Spyracopoulos L, LaPointe P (2014) A mutation in the catalytic loop of Hsp90 specifically impairs ATPase stimulation by Aha1p, but not Hch1p. J Mol Biol 426:2379–2392. PMID: 24726918 {Medline}

16. Hussein SK, Bhat R, Overduin M, LaPointe P (2024) Recruitment of Ahsa1 to Hsp90 is regulated by a conserved peptide that inhibits ATPase stimulation. EMBO Rep 25:3532–3546. PMID: 38937628 {Medline}

17. Jahn M, Rehn A, Pelz B, Hellenkamp B, Richter K, Rief M, Buchner J, Hugel T (2014) The charged linker of the molecular chaperone Hsp90 modulates domain contacts and biological function. Proc Natl Acad Sci U S A 111:17881–17886. PMID: 25468961 {Medline}

18. Knoblauch R, Garabedian MJ (1999) Role for Hsp90-associated cochaperone p23 in estrogen receptor signal transduction. Mol Cell Biol 19:3748–3759. PMID: 10207098 {Medline}

19. Koulov AV, LaPointe P, Lu B, Razvi A, Coppinger J, Dong MQ, Matteson J, Laister R, Arrowsmith C, Yates JR, 3rd, Balch WE (2010) Biological and structural basis for Aha1 regulation of Hsp90 ATPase activity in maintaining proteostasis in the human disease cystic fibrosis. Mol Biol Cell 21:871-884. PMID: 20089831 {Medline}

20. LaPointe P, Mercier R, Wolmarans A (2020) Aha-type co-chaperones: the alpha or the omega of the Hsp90 ATPase cycle? Biol Chem 401:423–434. PMID: 31782942 {Medline}

21. Lee CT, Graf C, Mayer FJ, Richter SM, Mayer MP (2012) Dynamics of the regulation of Hsp90 by the co-chaperone Sti1. EMBO J 31:1518–1528. PMID: 22354036 {Medline}

22. Li J, Richter K, Reinstein J, Buchner J (2013) Integration of the accelerator Aha1 in the Hsp90 co-chaperone cycle. Nat Struct Mol Biol 20:326–331. PMID: 23396352 {Medline}

23. Liu X, Wang Y (2022) Aha1 Is an Autonomous Chaperone for SULT1A1. Chem Res Toxicol 35:1418–1424. PMID: 35926086 {Medline}

24. Liu Y, Sun M, Myasnikov AG, Elnatan D, Delaeter N, Nguyenquang M, Agard DA (2020) Cryo-EM structures reveal a multistep mechanism of Hsp90 activation by co-chaperone Aha1. bioRxiv:2020.2006.2030.180695.

25. Lotz GP, Lin H, Harst A, Obermann WM (2003) Aha1 binds to the middle domain of Hsp90, contributes to client protein activation, and stimulates the ATPase activity of the molecular chaperone. J Biol Chem 278:17228–17235. PMID: 12604615 {Medline}

26. Mercier R, Wolmarans A, Schubert J, Neuweiler H, Johnson JL, LaPointe P (2019) The conserved NxNNWHW motif in Aha-type co-chaperones modulates the kinetics of Hsp90 ATPase stimulation. Nat Commun 10:1273. PMID: 30894538 {Medline}

27. Mercier R, Yama D, LaPointe P, Johnson JL (2023) Hsp90 mutants with distinct defects provide novel insights into cochaperone regulation of the folding cycle. PLoS Genet 19:e1010772. PMID: 37228112 {Medline}

28. Meyer P, Prodromou C, Liao C, Hu B, Roe SM, Vaughan CK, Vlasic I, Panaretou B, Piper PW, Pearl LH (2004) Structural basis for recruitment of the ATPase activator Aha1 to the Hsp90 chaperone machinery. EMBO J 23:1402–1410. PMID: 15039704 {Medline}

29. Mollapour M, Bourboulia D, Beebe K, Woodford MR, Polier S, Hoang A, Chelluri R, Li Y, Guo A, Lee MJ, Fotooh-Abadi E, Khan S, Prince T, Miyajima N, Yoshida S, Tsutsumi S, Xu W, Panaretou B, Stetler-Stevenson WG, Bratslavsky G, Trepel JB, Prodromou C, Neckers L (2014) Asymmetric Hsp90 N domain SUMOylation recruits Aha1 and ATP-competitive inhibitors. Mol Cell 53:317–329. PMID: 24462205 {Medline}

30. Mollapour M, Tsutsumi S, Donnelly AC, Beebe K, Tokita MJ, Lee MJ, Lee S, Morra G, Bourboulia D, Scroggins BT, Colombo G, Blagg BS, Panaretou B, Stetler-Stevenson WG, Trepel JB, Piper PW, Prodromou C, Pearl LH, Neckers L (2010) Swe1Wee1-dependent tyrosine phosphorylation of Hsp90 regulates distinct facets of chaperone function. Mol Cell 37:333–343. PMID: 20159553 {Medline}

31. Mollapour M, Tsutsumi S, Kim YS, Trepel J, Neckers L (2011) Casein kinase 2 phosphorylation of Hsp90 threonine 22 modulates chaperone function and drug sensitivity. Oncotarget 2:407–417. PMID: 21576760 {Medline}

32. Mollapour M, Tsutsumi S, Truman AW, Xu W, Vaughan CK, Beebe K, Konstantinova A, Vourganti S, Panaretou B, Piper PW, Trepel JB, Prodromou C, Pearl LH, Neckers L (2011) Threonine 22 phosphorylation attenuates Hsp90 interaction with cochaperones and affects its chaperone activity. Mol Cell 41:672–681. PMID: 21419342 {Medline}

33. Morishima Y, Murphy PJ, Li DP, Sanchez ER, Pratt WB (2000) Stepwise assembly of a glucocorticoid receptor.hsp90 heterocomplex resolves two sequential ATP-dependent events involving first hsp70 and then hsp90 in opening of the steroid binding pocket. J Biol Chem 275:18054–18060. PMID: 10764743 {Medline}

34. Obermann WM, Sondermann H, Russo AA, Pavletich NP, Hartl FU (1998) In vivo function of Hsp90 is dependent on ATP binding and ATP hydrolysis. J Cell Biol 143:901–910. PMID: 9817749 {Medline}

35. Panaretou B, Prodromou C, Roe SM, O’Brien R, Ladbury JE, Piper PW, Pearl LH (1998) ATP binding and hydrolysis are essential to the function of the Hsp90 molecular chaperone in vivo. EMBO J 17:4829–4836. PMID: 9707442 {Medline}

36. Panaretou B, Siligardi G, Meyer P, Maloney A, Sullivan JK, Singh S, Millson SH, Clarke PA, Naaby-Hansen S, Stein R, Cramer R, Mollapour M, Workman P, Piper PW, Pearl LH, Prodromou C (2002) Activation of the ATPase activity of hsp90 by the stress-regulated cochaperone aha1. Mol Cell 10:1307–1318. PMID: 12504007 {Medline}

37. Petropavlovskiy AA, Tauro MG, Lajoie P, Duennwald ML (2020) A Quantitative Imaging-Based Protocol for Yeast Growth and Survival on Agar Plates. STAR Protoc 1:100182. PMID: 33377076 {Medline}

38. Preuss KD, Pfreundschuh M, Weigert M, Fadle N, Regitz E, Kubuschok B (2015) Sumoylated HSP90 is a dominantly inherited plasma cell dyscrasias risk factor. J Clin Invest 125:316–323. PMID: 25485683 {Medline}

39. Prodromou C, Bjorklund DM (2022) Advances towards Understanding the Mechanism of Action of the Hsp90 Complex. Biomolecules 12. PMID: 35625528 {Medline}

40. Prodromou C, Siligardi G, O’Brien R, Woolfson DN, Regan L, Panaretou B, Ladbury JE, Piper PW, Pearl LH (1999) Regulation of Hsp90 ATPase activity by tetratricopeptide repeat (TPR)-domain co-chaperones. EMBO J 18:754–762. PMID: 9927435 {Medline}

41. Ratzke C, Mickler M, Hellenkamp B, Buchner J, Hugel T (2010) Dynamics of heat shock protein 90 C-terminal dimerization is an important part of its conformational cycle. Proc Natl Acad Sci U S A 107:16101–16106. PMID: 20736353 {Medline}

42. Reidy M, Garzillo K, Masison DC (2023) Nucleotide exchange is sufficient for Hsp90 functions in vivo. Nat Commun 14:2489. PMID: 37120429 {Medline}

43. Reidy M, Masison DC (2020) Mutations in the Hsp90 N Domain Identify a Site that Controls Dimer Opening and Expand Human Hsp90alpha Function in Yeast. J Mol Biol 432:4673–4689. PMID: 32565117 {Medline}

44. Retzlaff M, Hagn F, Mitschke L, Hessling M, Gugel F, Kessler H, Richter K, Buchner J (2010) Asymmetric activation of the hsp90 dimer by its cochaperone aha1. Mol Cell 37:344–354. PMID: 20159554 {Medline}

45. Richter K, Muschler P, Hainzl O, Buchner J (2001) Coordinated ATP hydrolysis by the Hsp90 dimer. J Biol Chem 276:33689–33696. PMID: 11441008 {Medline}

46. Richter K, Walter S, Buchner J (2004) The Co-chaperone Sba1 connects the ATPase reaction of Hsp90 to the progression of the chaperone cycle. J Mol Biol 342:1403–1413. PMID: 15364569 {Medline}

47. Siligardi G, Hu B, Panaretou B, Piper PW, Pearl LH, Prodromou C (2004) Co-chaperone regulation of conformational switching in the Hsp90 ATPase cycle. J Biol Chem 279:51989–51998. PMID: 15466438 {Medline}

48. Taipale M, Krykbaeva I, Koeva M, Kayatekin C, Westover KD, Karras GI, Lindquist S (2012) Quantitative analysis of HSP90-client interactions reveals principles of substrate recognition. Cell 150:987–1001. PMID: 22939624 {Medline}

49. Tang J, Hu H, Zhou C, Zhang N (2023) Human Aha1’s N-terminal extension confers it holdase activity in vitro. Protein Sci 32:e4735. PMID: 37486705 {Medline}

50. Tripathi V, Darnauer S, Hartwig NR, Obermann WM (2014) Aha1 can act as an autonomous chaperone to prevent aggregation of stressed proteins. J Biol Chem 289:36220–36228. PMID: 25378400 {Medline}

51. Van Oosten-Hawle P, Bolon DN, LaPointe P (2017) The diverse roles of Hsp90 and where to find them. Nat Struct Mol Biol 24:1–4. PMID: 28054566 {Medline}

52. Wandinger SK, Richter K, Buchner J (2008) The Hsp90 chaperone machinery. J Biol Chem 283:18473–18477. PMID: 18442971 {Medline}

53. Wang X, Venable J, LaPointe P, Hutt DM, Koulov AV, Coppinger J, Gurkan C, Kellner W, Matteson J, Plutner H, Riordan JR, Kelly JW, Yates JR, 3rd, Balch WE (2006) Hsp90 cochaperone Aha1 downregulation rescues misfolding of CFTR in cystic fibrosis. Cell 127:803–815. PMID: 17110338 {Medline}

54. Whitesell L, Mimnaugh EG, De Costa B, Myers CE, Neckers LM (1994) Inhibition of heat shock protein HSP90-pp60v-src heteroprotein complex formation by benzoquinone ansamycins: essential role for stress proteins in oncogenic transformation. Proc Natl Acad Sci U S A 91:8324–8328. PMID: 8078881 {Medline}

55. Wolmarans A, Kwantes A, LaPointe P (2019) A novel method for site-specific chemical SUMOylation: SUMOylation of Hsp90 modulates co-chaperone binding in vitro. Biol Chem 400:487–500. PMID: 30265648 {Medline}

56. Wolmarans A, Lee B, Spyracopoulos L, LaPointe P (2016) The Mechanism of Hsp90 ATPase Stimulation by Aha1. Sci Rep 6:33179. PMID: 27615124 {Medline}

57. Xu W, Mollapour M, Prodromou C, Wang S, Scroggins BT, Palchick Z, Beebe K, Siderius M, Lee MJ, Couvillon A, Trepel JB, Miyata Y, Matts R, Neckers L (2012) Dynamic tyrosine phosphorylation modulates cycling of the HSP90-P50(CDC37)-AHA1 chaperone machine. Mol Cell 47:434–443. PMID: 22727666 {Medline}

58. Zierer BK, Rubbelke M, Tippel F, Madl T, Schopf FH, Rutz DA, Richter K, Sattler M, Buchner J (2016) Importance of cycle timing for the function of the molecular chaperone Hsp90. Nat Struct Mol Biol 23:1020–1028. PMID: 27723736 {Medline}

